# Zinc and Metformin Co-Functionalized Polyetheretherketone: A Novel Dental Implant Material Tailored for the Elderly

**DOI:** 10.1101/2024.08.05.606613

**Authors:** Zhengwei Liu, Enze Zhao, Hanwei Huang, Yuxun Wu, Yicong He, Shuting Bai, Suwen Wang, Shirou Fan, Shuaishuai Cao, Bin Tang, Yansong Wang

**Author notes:** Corresponding author at : Department of Stomatology, Shenzhen Hospital (Futian) of Guangzhou University of Chinese Medicine, Shenzhen, Guangdong, P. R. China. These authors contributed equally to this work (Zhengwei Liu, Enze Zhao, Hanwei Huang).

## Abstract

This study focuses on addressing the challenges of dental implants in the geriatric population by enhancing the bioactivity of polyetheretherketone (PEEK) through surface modification. PEEK, with its elastic modulus close to alveolar bone, mitigates stress shielding but faces limitations in osseointegration due to low bioactivity. We introduced zinc (Zn) and metformin (MF) onto PEEK surfaces via a dopamine-assisted physical adhesion method, creating a functionalized derivative called ZnMF@PEEK. This combination targets diminished osteogenic potential, persistent inflammation, and cell senescence, which are common issues in elderly patients. Comprehensive physicochemical characterizations confirmed the successful preparation of ZnMF@PEEK, and in *vitro* and in *vivo* experiments systematically evaluated its biocompatibility and bioactivity. The results indicate that ZnMF@PEEK holds promise as a dental implant material tailored to the specific needs of the elderly, addressing multifaceted challenges in osseointegration.

## 1. Introduction

Dental implantology has made remarkable progress in recent decades, with titanium alloys becoming the standard material due to their excellent biocompatibility and mechanical strength[1]. However, a significant challenge with titanium alloys is their high elastic modulus, which surpasses that of human alveolar bone, leading to stress shielding[2, 3]. This issue can potentially reduce bone loading, causing bone resorption and increasing the risk of implant failure[4–6]. Alternatively, PEEK has gained attention as a potential material for future dental implants because its elastic modulus closely matches that of alveolar bone, reducing the risk of stress shielding and contributing to implant stability[7–9].

Despite PEEK’s favorable mechanical properties, its low bioactivity poses a challenge for achieving successful osseointegration, which is crucial for the long-term success of dental implants[10, 11]. Although PEEK-based materials with enhanced bioactivity have been developed for clinical use, they may not fully meet the unique needs of the geriatric population[12, 13]. The increasing demand for dental implants among older adults is further complicated by age-related factors such as reduced osteogenic capacity, persistent inflammation, and osteoblast senescence, all of which contribute to decreased bone formation efficiency[14–16]. Therefore, there is an urgent need to develop PEEK-based implant materials specifically tailored to the needs of older adults.

One scientifically validated and clinically feasible strategy to enhance PEEK’s osseointegration capabilities is surface modification with bioactive substances[17, 18]. Zinc (Zn), an essential trace element, is well-known for its role in stimulating osteoblast differentiation and mineralization, as well as its anti-inflammatory properties[19]. On the other hand, MF, a long-standing diabetes medication, has recently gained attention for its unique ability to combat cellular senescence and potentially promote osteogenesis[20]. The combination of Zn and MF holds promise for addressing the multifaceted challenges of osseointegration in older adults, specifically reduced osteogenic capacity, chronic inflammation, and osteoblast senescence[21, 22].

In this study, we utilized a dopamine-assisted physical adhesion method to introduce Zn and MF onto the surface of PEEK, resulting in a functionalized PEEK derivative called ZnMF@PEEK. We hypothesize that the combination of Zn and MF will effectively target the three main issues related to bone integration in older adults: reduced osteogenic potential, persistent inflammation, and osteoblast senescence. To validate the successful preparation of ZnMF@PEEK, we conducted a series of physicochemical characterizations. Additionally, we performed comprehensive in *vitro* and in *vivo* experiments to systematically evaluate its biocompatibility and bioactivity. The results indicate that ZnMF@PEEK is a promising candidate for dental implant materials in the geriatric population.

## 2. Experimental details

### 2.1 Materials

MF was purchased from Bio-Year Technology Co., Ltd. (China), PEEK was purchased from Victrex Technology Company (UK), and dopamine, NaOH, ZnCl2 and cation exchange resin were all purchased from Sigma-Aldrich Company (China). All reactants were used as received. Deionized water (> 18.2 MΩ·cm) was used when water was involved.

### 2.2 Preparation of ZnMF@PEEK

Zinc metformin (ZnMF) was prepared beforehand. First, a saturated aqueous solution of MF was mixed with an equal volume of 2M aqueous ZnCl2 solution. The mixed solution was then stirred with NaOH to adjust the pH to be in the range of 9 to 11 for 1 h to ensure complete reaction. Subsequently, absolute ethanol was added to the reaction mixture and left for 24 h to allow full precipitation of the composite powder. The obtained composite powder was filtered using a vacuum suction device. After filtration, it was stirred and washed with ethanol aqueous solutions at concentrations of 80%, 60%, 40%, respectively. To ensure that the residual ZnCl2 was completely removed, the washing procedure was repeated three times, and the final product after the washing was ZnMF.

Medical-grade PEEK samples were washed with acetone, absolute ethanol, and deionized water under ultrasound for 10 min, and then dried in an incubator at 60 °C for 24 h. Afterwards, the PEEK samples were sulfonated by immersing them in 98% concentrated sulfuric acid and ultrasonically vibrating them at 25°C for 5 minutes. After sulfonation, the PEEK samples were cleaned under ultrasonic conditions with a ratio of 5 ml of deionized water for 10 minutes twice to remove residuals. After cleaning, they were dried in a constant temperature oven at 60 °C for 24 h. The dried PEEK sample was named SPEEK.

An aqueous dopamine solution was prepared by dissolving dopamine in Tris buffer at a concentration of 2 mg/mL and pH 8.5. The SPEEKs were soaked in the aqueous dopamine solution for 24 h to introduce dopamine onto the SPEEK surface, and then rinsed with deionized water to remove any unattached residues. The SPEEK after the introduction of dopamine was thereafter named DSPEEK. we administered 200 μL of ZnMF solution onto the surface of DSPEEK and allowed it to remain at room temperature for a duration of 24 hours, ensuring that ZnMF was fully bound to DSPEEK. This process resulted in the formation of PEEK with incorporated ZnMF, which we designated as ZnMF@PEEK. Subsequently, these samples underwent freeze-drying for a period of 48 hours. To investigate the effects of varying ZnMF concentrations, we prepared a series of ZnMF@PEEK samples using ZnMF solutions with concentrations of 100, 300, 500, 1000, and 1500 μg/mL.

### 2.3 Materials characterizations

The microstructure of the samples was observed using a scanning electron microscope (JSM- 7200F, JEOL company, Japan). The surface microstructure of PEEK samples and prepared samples was observed by scanning electron microscopy (SEM, Nova NanoSem450); the elemental composition and element distribution of the surface of different samples were measured by energy dispersive spectrometer (EDS) (Nova NanoSem450 FEI, USA). Since the samples are non- conductive, prior to SEM observation, gold spraying was performed to enhance conductivity and optimize image clarity. In addition to EDS, attenuated total reflection Fourier transform infrared spectroscopy (ATR-FTIR) (VERTEX 70v, German) and Ultraviolet–visible spectroscopy (UV- Vis) were employed to further investigate the chemical structure (UV-2600 SHIMADZU, Japan).

The Zn ion release behavior of the samples was investigated using inductively coupled plasma- optical emission spectrometry (ICP-MS) (Agilent 7900 ICP-MS, USA). Samples were soaked in physiological saline at 37°C to simulate the in *vivo* environment. Aliquots were collected at pre-designed time points over a 72-hour period, and the concentration of Zn ions in each aliquot was measured respectively.

The surface hydrophilicity of the samples was quantitatively assessed using a contact angle system (VCA OPTIMA, AST) In the contact angle measurement, three samples from each group were used, with measurements taken at five randomly selected points on each sample to improve the reliability of the results.

### 2.4 Cell culture

Mouse embryonic osteoblasts MC3T3-E1 cells were used in this study. The culture medium used was α-MEM containing 1% penicillin-streptomycin and 10% FBS, and the cells were cultured in a cell incubator under standard culture conditions (37°C, 5% CO2). The culture medium was regularly replaced every 2 days to ensure healthy cell growth and maintain their differentiation potential. After 5 days, cells were passaged to avoid contact inhibit.

### 2.5 Cytotoxicity assay

In this study, the CCK8 method was used to evaluate the cell viability of samples and to determine the optimal drug concentration. During the experiment, MC3T3-E1 cells were first evenly seeded in a 96-well plate at a density of 1×104 cells/well, and an appropriate amount of culture medium was added. Incubate under constant temperature culture conditions of 37°C and 5% CO2. After one day of cell attachment growth, the original medium was replaced with culture medium containing different sample concentrations, including control group with PEEK (NC), DSPEEK group (0), and ZnMF@PEEK synthesized at 100, 300, 500, 1000, and 1500 μg/ml. Each sample was immersed in 15 ml of complete medium and shaken in a shaker at 37 °C for 24 h to get sample effusion. Subsequently, at each set time node, 10 μL of CCK8 solution with concentration of 5 mg/mL was added to each well, and incubation continued for 4 h. After the incubation, carefully remove the culture medium and detect the absorbance value of each well at a wavelength of 450 nm using a microplate reader. To ensure the reliability of the experimental results, each set of experiments was repeated three times. By comparing cell viability at different sample concentrations, the experimental group can evaluate the cytotoxicity of the sample and determine the optimal drug concentration accordingly.

The dead/live cell staining method was used to further assess the cell viability and possible cell cytotoxicity. MC3T3-E1 cells with initial density 2×104 cells/well were first seeded on the surface of the material in the well plate of culture medium. Subsequently, the plates were placed in a cell incubator at 37°C and 5% CO2 for 1, 3, and 5 days, respectively. Calcein-AM/PI Double Straining Kit (Beyotime, China) was used for the staining, and an inverted fluorescence microscope was employed to observe stained cells.

### 2.6 Real-time fluorescence quantitative PCR(RT-qPCR)

qPCR was used to analyze the expression of bone-promoting genes, inflammatory genes before and after implant modification. (1) Inoculate MC3T3-E1 at a ratio of 1 x 105 cells/well on sterilized DSPEEK placed in a 6-well plate and culture for 7 days. (2) Use a pipette to precipitate the culture medium in the well and clean it with PBS. Add 1 ml of Trizol RNA extraction solution, mix with a pipette repeatedly, then send it to a 2 ml centrifuge, add 200 μl of chloroform, shake and mix for 20 s, and then centrifuge at 12000 rpm at 4°C. 15 minutes. (3) The solution after centrifugation is divided into three layers by naked eye: the top layer is the aqueous phase, the second layer is the protein, and the third layer is the organic phase. Gently take out 500 μl of the supernatant without touching the protein and organic phase. After taking out the aqueous phase solution, add 500 μl of the same amount of isopropyl alcohol solution. After mixing, rotate at: 12000 rpm; temperature: 4°C; Time: centrifuge for 15 minutes. (4) After the pre-centrifugation process, white precipitate accumulated at the bottom of the centrifuge tube with the naked eye. Aspirate the solution, add absolute ethanol solution, and wash the white precipitate. (5) Use DEPC water to dissolve the white precipitate at the bottom of the centrifuge tube, and place it in the instrument to detect the concentration and purity of the extracted RNA. (6) Prepare 30 μl of reverse transcription solution according to the instructions of the purchased reverse transcription kit manufacturer, in which 1 μl of RNA, 4 μl of 5X Evo-M-MLVRT master Mix, and 15 μl of sterile enzyme-free water are added. After it is ready, place it in the PCR instrument to perform the reverse transcription reaction. (7) Finally, the experiment was normalized by glyceraldehyde- 3-phosphate dehydrogenase (GADPH), and gene expression was calculated according to the 2− ΔΔCt method [17]. The primers used are shown in Table 1.

**Table 1.**
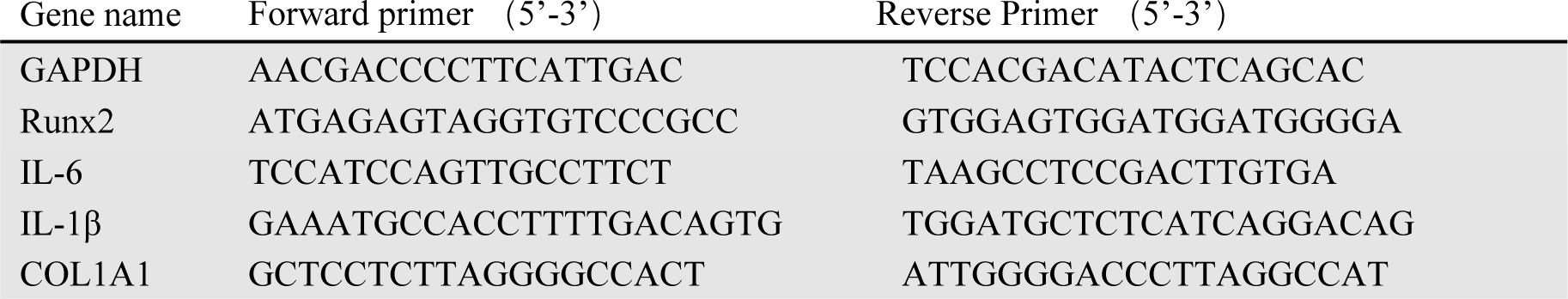
The sequence of the primer for RT-PCR.

### 2.7 Alizarin red staining and alkaline phosphatase (ALP) immunofluorescence staining

After culturing MC3T3-E1 cells on various PEEK material for 7 days, the cells were delicately washed three times with PBS to eliminate any possible contaminants and unattached cells. The cells were then stabilized using 4% paraformaldehyde for 15 minutes at ambient temperature to preserve their morphological structure. Following this, the cells underwent another triple wash with deionized water to rid of any leftover fixing agent. After that, the cells were stained with alizarin red kit supplied by Beyotime company in China, following the instruction provided by the supplier. Upon completion of staining, the alizarin red solution was discarded, and the samples were rinsed with deionized water until all unbound dye was removed.

Immunofluorescence (IF) staining was conducted to detect alkaline phosphatase (ALP). Following a ten-day incubation period, the culture medium was carefully aspirated, and MC3T3-E1 cells were rinsed twice with PBS. The cells were then immobilized for 15 minutes at ambient temperature using Immunol Staining Fix Solution (P0098, Beyotime) before undergoing three washes with Immunol Staining Wash Buffer (P0106, Beyotime). The MC3T3-E1 cells were subsequently permeabilized and blocked with Immunol Staining Blocking Buffer (P0102, Beyotime) for one hour. The primary antibody targeting ALP was applied to the MC3T3-E1 cells and incubated at 4°C throughout the night. After three washes with Immunol Staining Wash Buffer, the samples were labeled with fluorescein isothiocyanate (FITC) serving as the secondary antibody. For nuclear staining, the samples were exposed to a diluted solution of Hoechst 333258 dye for five minutes at room temperature. To assess ALP synthesis, confocal microscopy was employed. The microscopic images of the MC3T3-E1 cells were captured using a confocal laser scanning microscope (CLSM, TCS SP8, Leica, Germany).

### 2.8 Animal model

To investigate dental implant integration and healing performance, 16 young Sprague Dawley (SD) male rats, aged 8-10 weeks and weighing 200-250g, were used to create a femoral defect model to mimic the osteointegration of dental implant. The animals are randomly assigned to four groups: PEEK, SPEEK, DSPEEK, and ZnMF@PEEK.

After anesthetizing the rats with isoflurane inhalation, a uniformly sized and positioned bone defect (ø2 mm × 3 mm × 3 mm) was surgically created on the right femur to mimic the environment for a dental implant. The defect was then cleaned with physiological saline and filled with PEEK samples matching its dimensions. After the implantation, the incision at the surgical site was sutured and disinfected with iodophor. To prevent infection, all the animals were intraperitoneally injected with penicillin (100,000 units kg^−1^) after the surgery.

### 2.9 Micro-CT investigation

All animals in the experimental group underwent euthanasia after a two-week recovery period. Following this, an incision was made to remove the skin and muscle tissue, allowing for the extraction of the bone defect specimens. A Micro-CT((Kontich, Belgium) was employed to scan the specimens and perform the quantitative analysis to obtain the bone structural parameters, including bone mineral density(BMD) and bone volume fraction of trabeculae(BV/TV). This comprehensive assessment facilitated a comparison of the osseointegration status across different groups.

### 2.10 Tissue staining

To observe the bone-implant integration, The standard Masson and hematoxylin-eosin (HE) staining techniques were employed. Prior to staining, the bone samples were thoroughly cleansed with ethanol. The specimens then underwent a series of processes, including dehydration, infiltration, and embedding, before being cut and ground into tissue sections. These sections were subsequently stained using Masson and HE techniques. Furthermore, key organ tissues, including the heart, liver, spleen, lungs, and kidneys, were also collected postmortem from the sacrificed animals. These tissues were subjected to HE staining to assess the long-term safety of the implant. Histological analysis were performed to evaluate the ZnMF@PEEK’s potential in anti-osteoblast aging. Immunohistochemistry staining of P53, P21 and β-galactosidase were used to assess the aging of anti-osteoblast. Slices sliced in succession were subjected to sodium citrate buffer (pH 6.0) at 58 ℃ for 16 h for antigen retrieval and then incubated with rabbit anti-P53 and P21 antibodies, respectively. For β-galactosidase staining, slides were subjected to sodium citrate buffer (pH 6.0) at 58 ℃ for 16 h for antigen retrieval. And then incubated with rabbit anti-β- galactosidase antibodies after HE staining. Visualization was achieved using DAB substrate, and hematoxylin was used for counterstaining. The prepared slides were then dehydrated, cleared, and mounted for microscopic examination.

### 2.11 Statistical analysis

All the images analysis were processed and analyzed with Image J. The statistical significant difference in experiment data was analyzed with Student’s t-test. All values involved in the study were denoted as mean ± standard deviation (SD). *p* < 0.05 was regarded as the appearance of a significant difference in the data.

## 3. Results

### 3.1 Materials characterizations of various PEEK samples

In Figure 1, the SEM morphology (Figure 1A-F) and EDS surface element analysis results (Figure 1G-J) of various samples are presented. The SEM images (Figure 1A) reveal that MF exhibits a regular prismatic crystal structure, whereas ZnMF (Figure 1B) demonstrates a significant transformation into a disordered granular structure. PEEK (Figure 1C) shows a relatively smooth surface, but upon sulfonation, SPEEK (Figure 1D) acquires a prominent three-dimensional mesh-like porous structure with pore sizes ranging from 0.5 to 5 μm. Further processing to obtain DSPEEK (Figure 1E) does not alter its microstructure or pore size significantly. Notably, ZnMF@PEEK (Figure 1F) maintains a similar microstructure to DSPEEK, with a slight reduction in pore size, primarily distributed between 0.5 and 2 μm, and distinctive granular objects evenly dispersed on its surface. Additionally, EDS analysis confirms the absence of zinc on the surfaces of PEEK (Figure 1G), SPEEK (Figure 1H), and DSPEEK (Figure 1I), while a considerable amount of zinc is detected on the surface of ZnMF@PEEK (Figure 1J), indicating the successful incorporation of ZnMF on DSPEEK surface.

**Figure 1.**
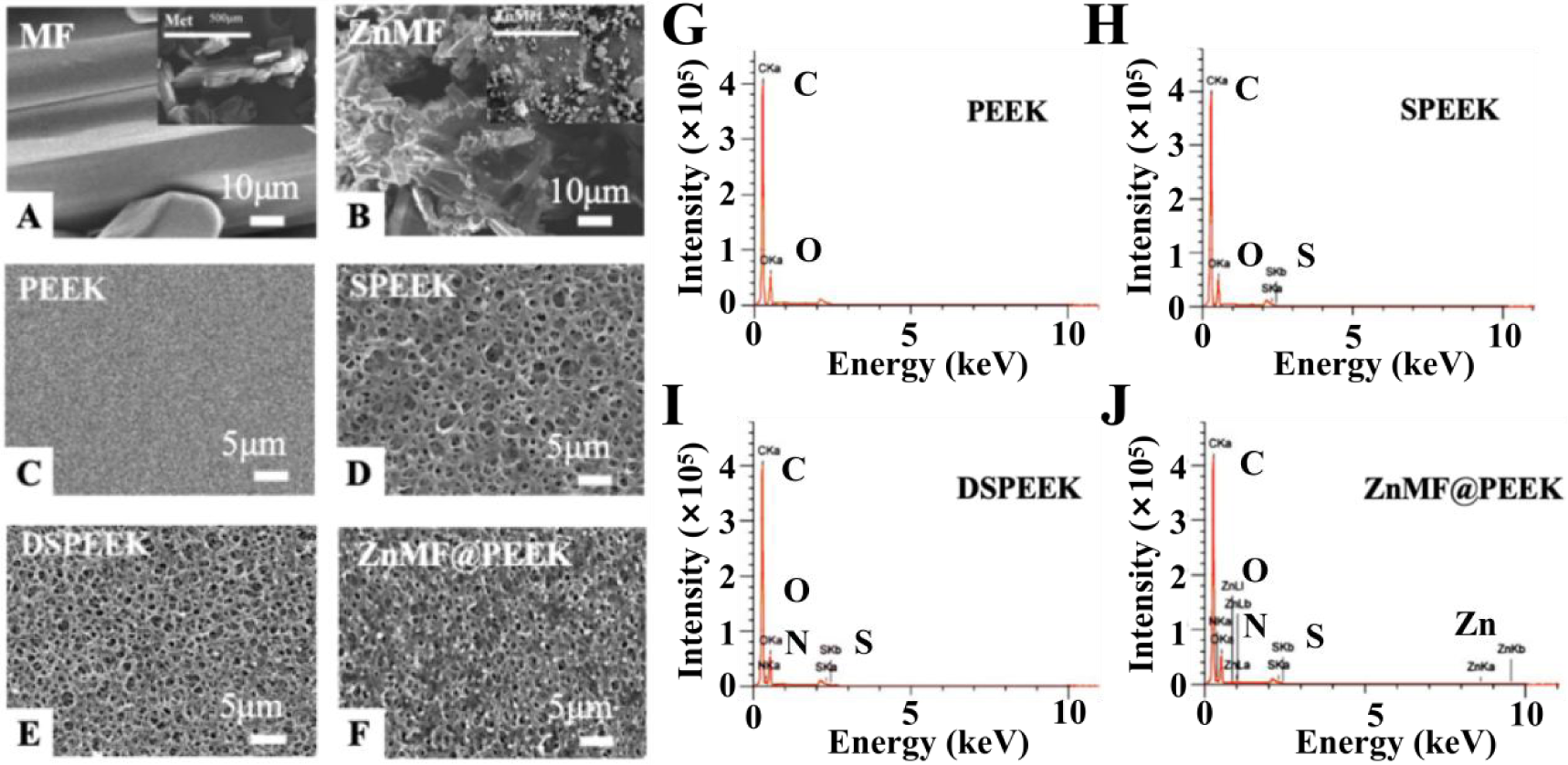
SEM images and EDS analysis of various samples. (A) MF displaying a regular prismatic crystal structure. (B) ZnMF showing a transition to a disordered granular structure. (C) Smooth surface of PEEK. (D) SPEEK exhibiting a distinct three-dimensional porous network with pores ranging from 0.5 to 5 μm. (E) DSPEEK maintaining a similar microstructure to SPEEK. (F) ZnMF@PEEK with pores primarily in the 0.5 to 2 μm range and ZnMF nanoparticles evenly distributed on the surface. (G-J) EDS spectra indicating the absence of zinc on PEEK (G), SPEEK (H), and DSPEEK (I) surfaces, whereas a significant amount of zinc is detected on the ZnMF@PEEK surface (J).

The UV-Vis absorption spectra presented in Figure 2A depict the optical properties of MF and ZnMF. Both MF and ZnMF exhibit distinct absorption peaks centered around 255 nm, indicative of electronic transitions within their respective molecular structures. Notably, the absorption peak of ZnMF is observed to be more pronounced and exhibits a higher absorbance intensity compared to that of MF. This observation suggests an enhancement in the UV absorption properties of MF upon zinc complexation, potentially attributed to the altered electronic configuration and/or increased conjugation within the ZnMF molecule[23, 24].

**Figure 2.**
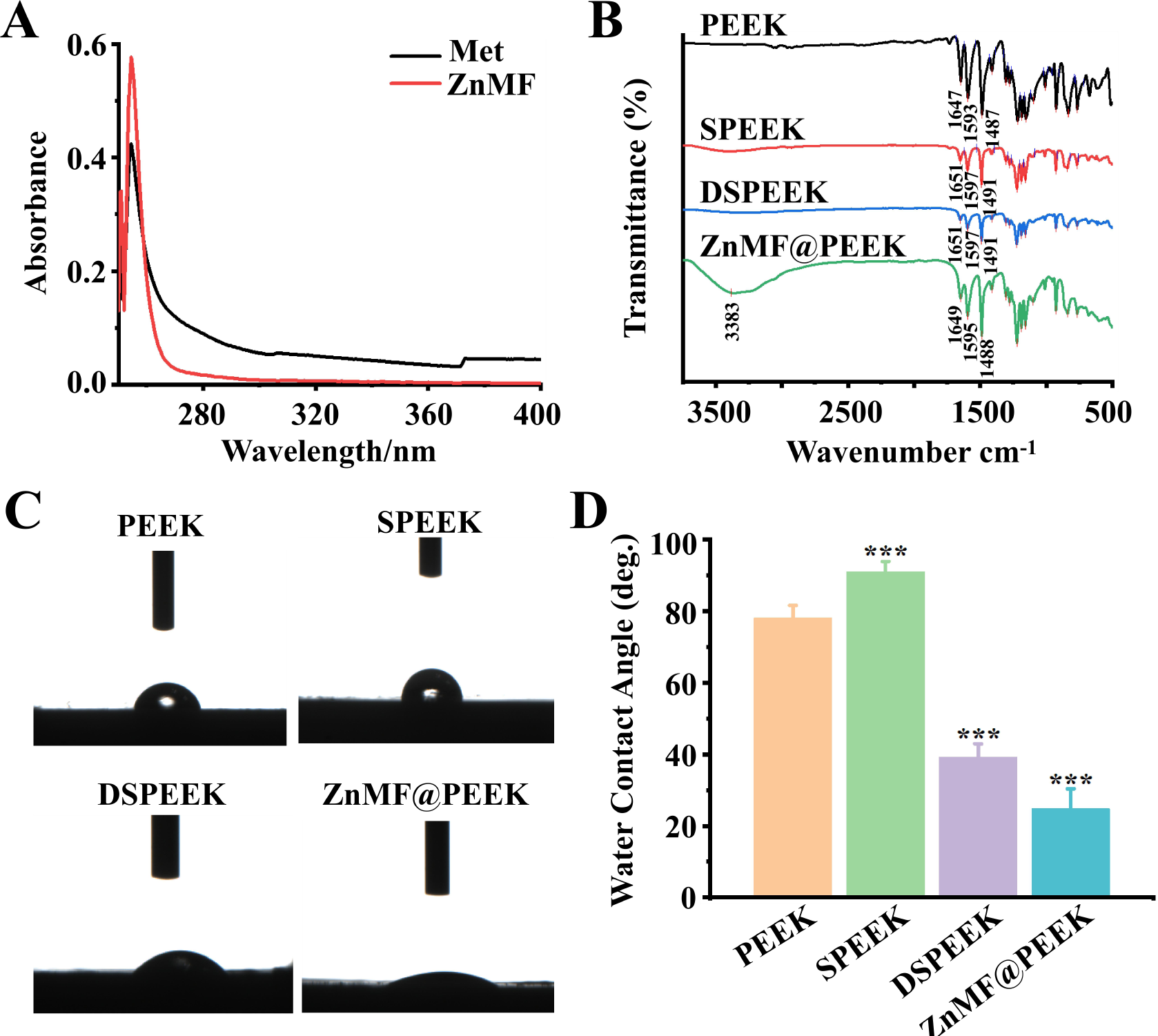
Materials characterization and wettability results of various samples. (A) UV-Vis absorption spectra of MF and ZnMF. (B) FTIR spectra of PEEK, SPEEK, DSPEEK, and ZnMF@PEEK. (C) Optical photographs of contact angle test images for different PEEK samples. (D) Quantitative contact angle data for the PEEK samples.

The FTIR spectra of PEEK, SPEEK, DSPEEK, and ZnMF@PEEK presented in Figure 2B reveal distinct absorption features characteristic of their chemical structures. PEEK exhibits prominent peaks at 1647, 1593, and 1487 cm^-1^, attributed to the aromatic backbone vibrations[25, 26]. SPEEK shares similar spectral features with PEEK but exhibits an additional peak at 1051 cm^-1^, indicative of the successful sulfonation process and the presence of sulfonic acid groups[27]. DSPEEK retains the spectral profile of SPEEK, suggesting minimal changes to the bulk structure upon dopamine modification. Notably, ZnMF@PEEK displays a novel peak at 3364 cm^-1^, which is due to the N-H symmetric stretching vibrations, should arise from the introduction of ZnMF[28]. Figure 2C presents optical photographs showing the contact angle test images for different PEEK samples, which clearly showcase the wettability characteristics of each material, allowing a direct visual comparison among them. Complementing these qualitative observations, Figure 2D quantitatively outlines the contact angle data. The contact angle for pure PEEK is smaller than that of SPEEK. However, when compared to DSPEEK, PEEK displays a larger contact angle, suggesting relatively poorer wettability[29]. Remarkably, ZnMF@PEEK demonstrates the smallest contact angle in the series, indicating the most enhanced wettability among all the tested samples.

The results presented in Figure 3 show the cumulative Zn ion release profile of the ZnMF@PEEK sample over a 24-hour period. It was observed that Zn ions were rapidly released within the first 8 hours, followed by a gradual slowing down of the release rate, reaching a plateau phase after approximately 12 hours.

**Figure 3.**
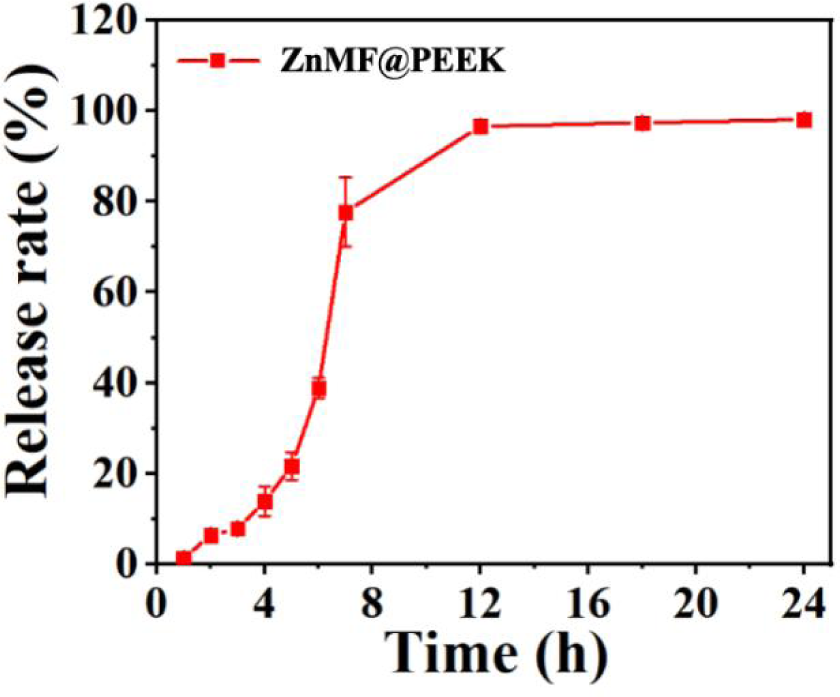
Cumulative Zn ion release curve of ZnMF@PEEK measured with ICP-MS.

### 3.2 In *vitro* assessment the bioactivities of various PEEK samples

The cell viability and potential toxicity of various PEEK samples were investigated using MC3T3- E1 cells, with DSPEEK serving as the control group, as shown in Figure 4. Figure 4A presents the cell viability data of ZnMF@PEEK samples prepared with different ZnMF concentrations. Consistent with the control group, the absorbance values, which reflect the count of living cell number, for all ZnMF@PEEK samples significantly increased over time, preliminarily indicating the safety of ZnMF@PEEK. Regarding the absorbance values across groups, no statistically significant differences were observed on days 1 and 3. However, on day 5, the absorbance values for the 500 μg/ml and 1000 μg/ml groups were significantly higher than those of the control group. Although there was no statistical difference in absorbance between the 500 μg/ml and 1000 μg/ml groups, the absorbance value was higher in the 500 μg/ml group. Therefore, this concentration was selected for further exploration of material safety through live/dead staining experiments. Figure 4B shows typical live/dead staining images of PEEK, DSPEEK, and ZnMF@PEEK (500 μg/ml) samples. Notably, no significant dead cells were observed in any group at each time point, and the living cell increased markedly over time in all groups. Compared to PEEK and DSPEEK, the ZnMF@PEEK samples exhibited a more pronounced increase in cell proliferation at all time points. The results shown in Figure 4 indicate that ZnMF@PEEK at 500 μg/ml exhibits enhanced cell viability. Therefore, this specific ZnMF@PEEK group was uniformly adopted for subsequent experiments of bioactivities assessment.

**Figure 4.**
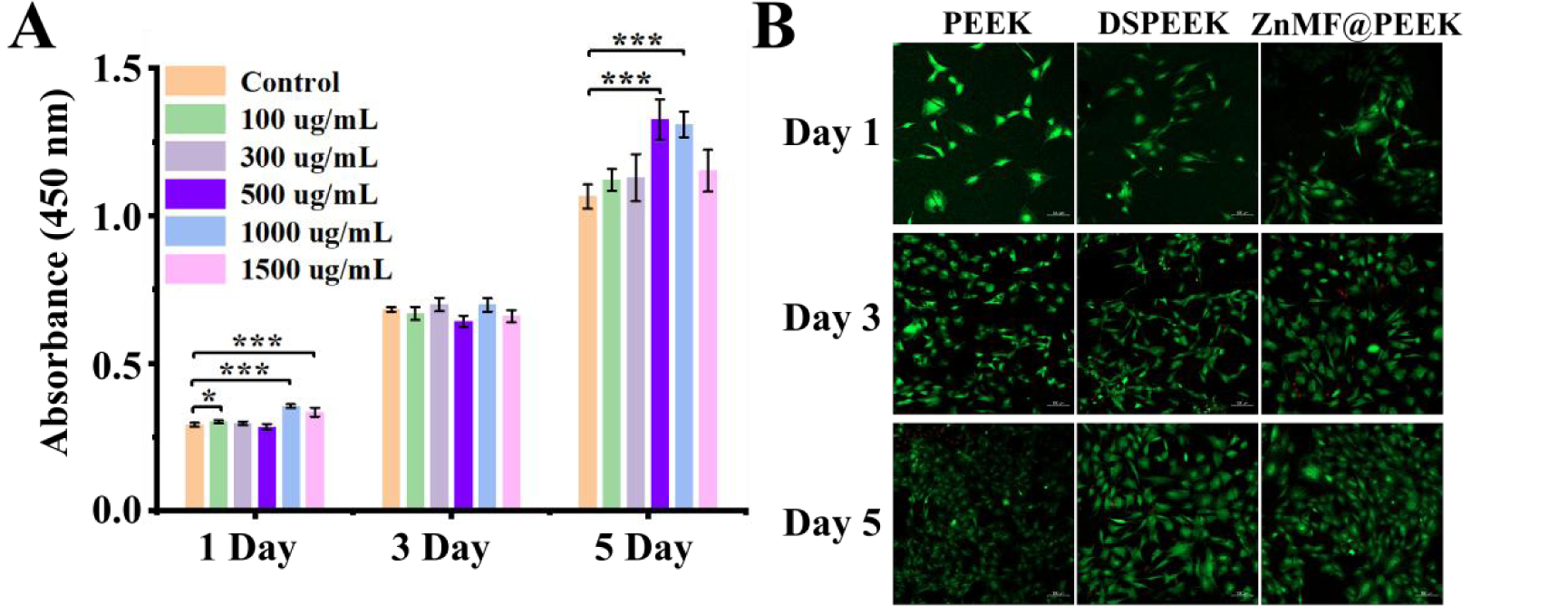
Cell Viability and Safety Assessment of ZnMF@PEEK Samples. (A) Cell viability data of ZnMF@PEEK samples prepared with different ZnMF concentrations. (B) Live/dead staining images of PEEK, DSPEEK, and ZnMF@PEEK (500 μg/ml) samples at different time points.

The regulatory effects of various PEEK materials on osteoblastogenesis and anti-inflammatory related genes in MC3T3-E1 cells was investigated, as shown in Figure 5. Here, conventional culture dishes were employed as the control group for comparative analysis.

**Figure 5.**
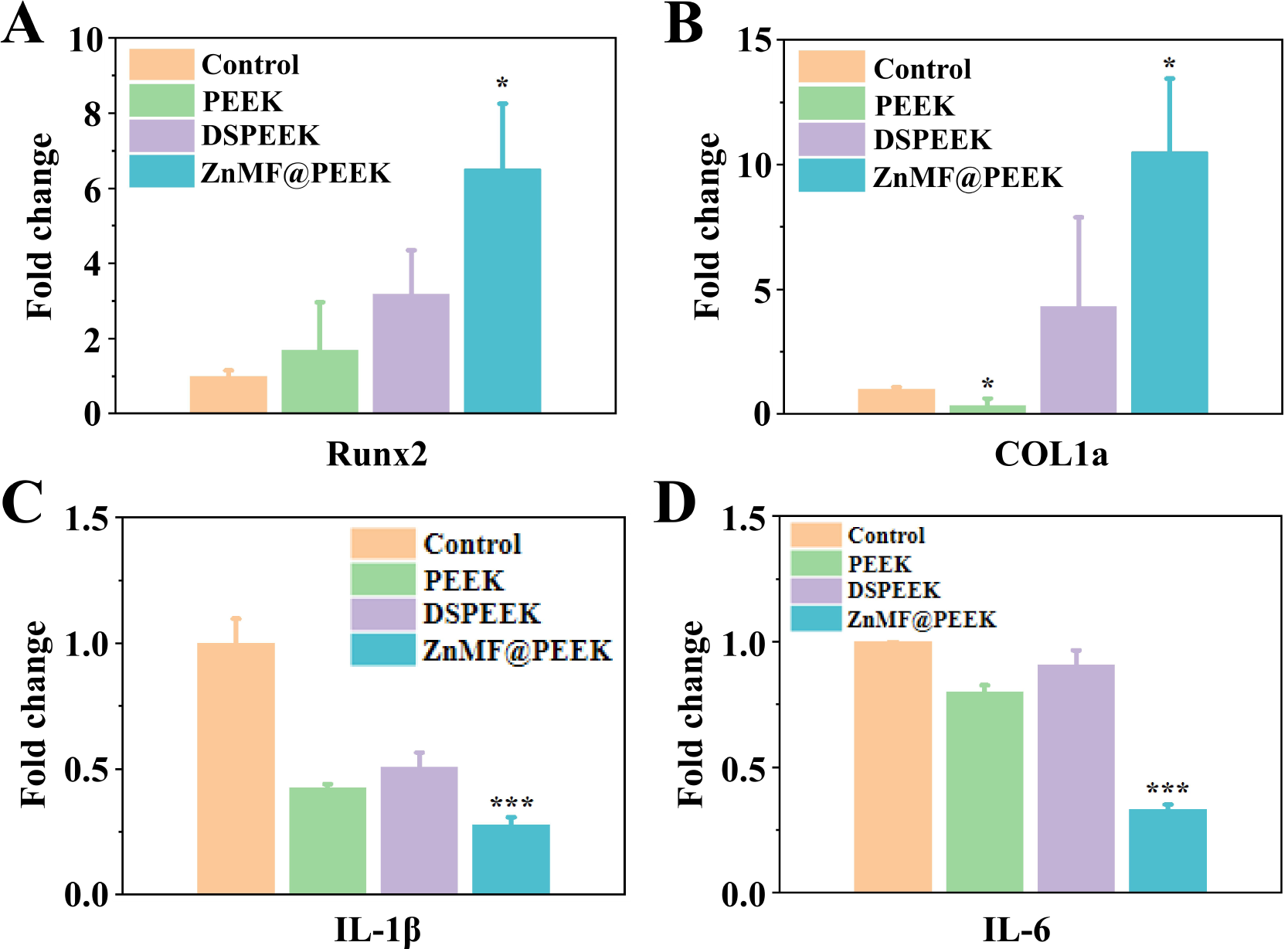
Regulatory effects of various samples on osteogenesis and anti-inflammatory related genes in MC3T3-E1 cells, (A) Runx2, (B) COL1a, (C) IL-1b and (D)IL-6.

Regarding the expression of RUNX2 (Figure 5A), compared to the control group, PEEK material significantly upregulated its expression level. Notably, the modified DSPEEK group demonstrated even higher RUNX2 expression than the PEEK group, further enhancing the potential for osteogenic differentiation. It is worth mentioning that the ZnMF@PEEK group exhibited the most significant increase in RUNX2 expression, suggesting that ZnMF@PEEK may possess the optimal ability to promote osteogenic differentiation[30, 31]. In contrast to the RUNX2 expression pattern, we observed a different trend in COL1a expression (Figure 5B). Compared to the control group, PEEK material significantly downregulated COL1a expression. However, the DSPEEK group showed a clear upregulation of COL1a expression compared to the control group, indicating its unique effect in promoting osteogenic differentiation[32, 33]. Remarkably, the ZnMF@PEEK group demonstrated the most prominent upregulation of COL1a, significantly higher than the DSPEEK group, further confirming its advantage in osteogenic differentiation.

Regarding the inflammation-related genes IL-1β (Figure 5C) and IL-6 (Figure 5D), we also conducted detailed examinations. The results revealed that, in terms of IL-1β expression, all PEEK groups exhibited significantly lower expression levels compared to the control group. Notably, the ZnMF@PEEK group demonstrated the most significant downregulation, suggesting that ZnMF@PEEK may possess the strongest anti-inflammatory effect[34]. Similarly, compared to the control group, PEEK also showed a significant downregulation trend in IL-6 expression. Although the DSPEEK group had slightly higher expression than the PEEK group, it was still lower than the control group. It is noteworthy that the ZnMF@PEEK group exhibited the lowest expression of IL-6, further confirming its significant anti-inflammatory effect.

The osteogenic differentiation and bone mineralization capabilities of MC3T3-E1 cells on various material surfaces were evaluated, with the control being the standard cell culture dish surface. Figure 6A presents the immunofluorescence staining images of ALP after 5 days of culturing on different sample surfaces. Notably, the ZnMF@PEEK group exhibited higher ALP expression compared to the other groups. This observation was corroborated by the quantitative analysis based on the staining images shown in Figure 6B, where the ZnMF@PEEK group demonstrated significantly higher fluorescence intensity than the rest of the groups. Interestingly, the PEEK group also showed a marked increase in ALP fluorescence expression compared to the control, indicating its potential to enhance osteogenic differentiation[35, 36]. Furthermore, Figure 6C displays the Alizarin red staining images, which revealed more pronounced calcified nodules in the ZnMF@PEEK group, suggesting its superior ability to promote bone mineralization.

**Figure 6:**
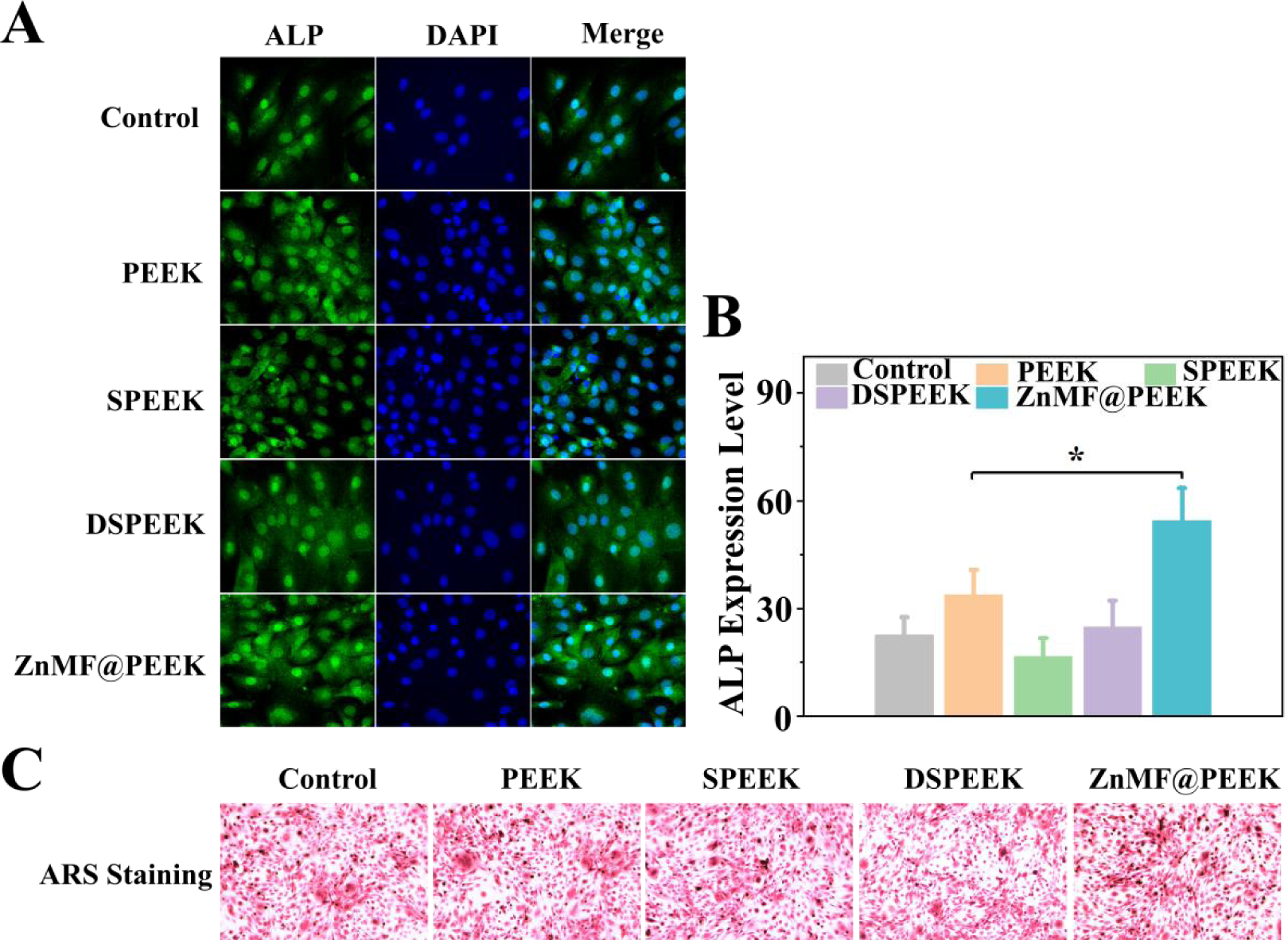
Assessment of osteogenic differentiation and bone mineralization capabilities of MC3T3-E1 cells on different material surfaces. The control represents the standard tissue culture plate surface. (A) Immunofluorescence staining images of ALP (alkaline phosphatase) after 5 days of culturing, (B) Quantitative analysis of ALP fluorescence intensity, (C) Alizarin red staining images.

### 3.2 In *vivo* Assessment of ZnMF@PEEK for Orthopedic Implants

To assess the osseointegration capabilities of these dental implant materials, we employed a femoral defect model following standard protocols established in previous studies. The preliminary animal experimental outcomes evaluating the bone integration of PEEK and ZnMF@PEEK are presented in Figure 7. The construction process of the animal model is illustrated in Figure 7A. 4 and 8 weeks post-implantation, femoral tissue samples containing the implants were harvested. Initial gross examination revealed no signs of infection in any of the tissues, with ZnMF@PEEK appearing superior bone integration compared to PEEK.

**Figure 7:**
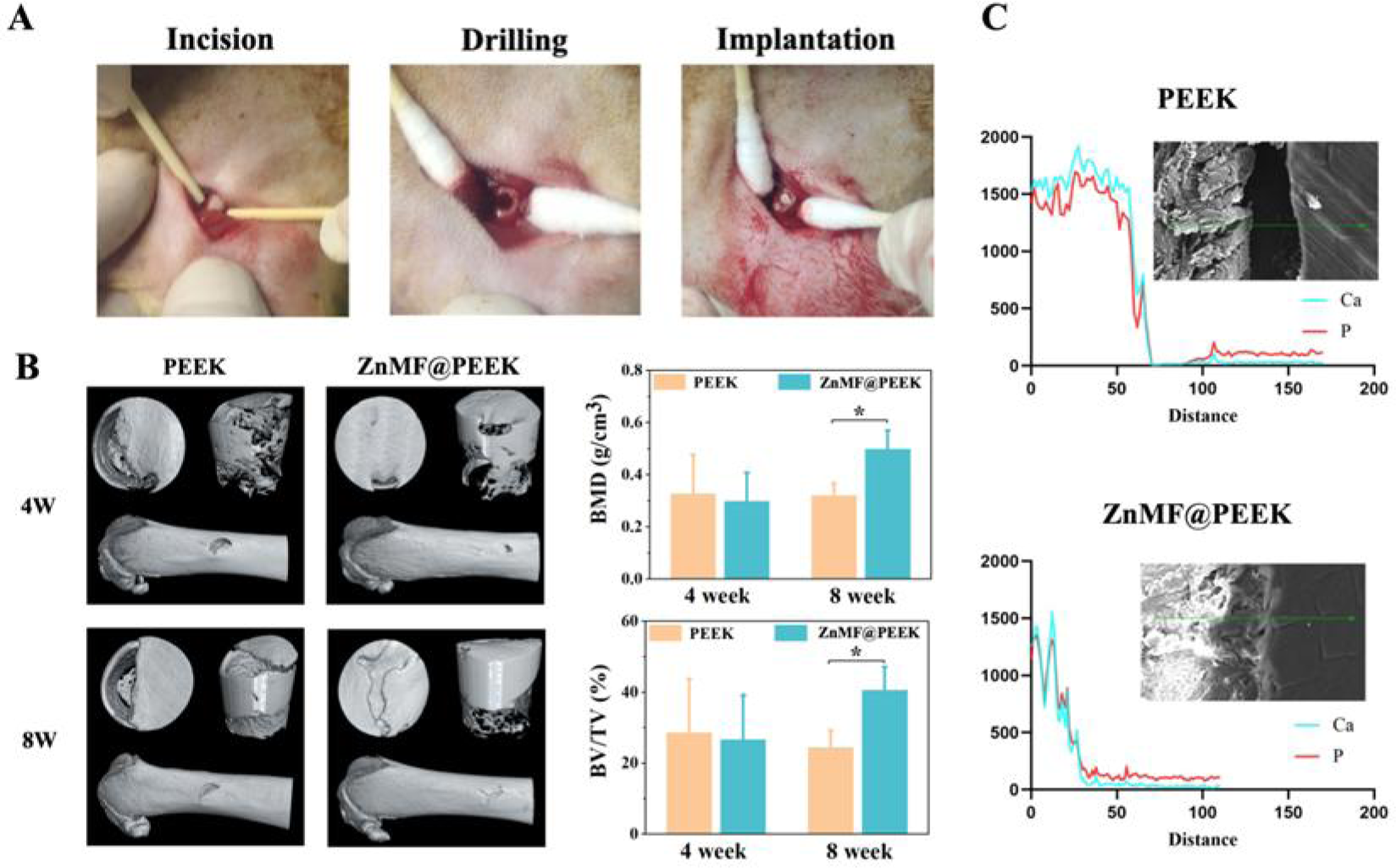
Assessment of Osseointegration Capabilities of PEEK and ZnMF@PEEK Dental Implant Materials. (A) Illustration of the femoral defect model construction process used in the animal experiments, (B) Micro-CT scanning images comparing bone formation around PEEK and ZnMF@PEEK implants at 4 and 8 weeks post-implantation, (C) EDS line scan results across the implant-bone interface for both PEEK and ZnMF@PEEK.

To validate these visual observations, micro-CT scanning analysis and Energy-Dispersive Spectroscopy (EDS) line scans across the implant-bone interface were performed. The micro-CT images presented in Figure 7B confirm the visual assessment, showing a higher amount of new bone formation around ZnMF@PEEK compared to PEEK, particularly evident at 8 weeks post-implantation. This observation is further supported by quantitative analyses based on CT scan data, which indicate significantly higher BMD and BV/TV values around ZnMF@PEEK implants at 8 weeks compared to PEEK implants.

Furthermore, EDS line scan results presented in Figure 7C reveal distinct differences in the distribution of calcium and phosphorus elements across the implant-bone interface between the two groups. In the PEEK group, a sharp decrease in calcium and phosphorus concentrations is observed as the scan crosses the interface, indicating limited bone integration. In contrast, the ZnMF@PEEK group shows a more gradual decline in these element concentrations, suggesting better integration between the modified implant and bone tissue.

The histological analysis presented in Figure 8 provides further evidence of better osteointegration capabilities of ZnMF@PEEK, as compared to the PEEK group. Masson staining (Figure 8A) reveals a notably higher distribution of collagen fibers for ZnMF@PEEK samples. These collagen fibers are not only more abundant but also exhibit a uniform arrangement, indicative of improved bone formation and matrix organization. This observation suggests that the ZnMF@PEEK composite fosters a more conducive environment for collagen deposition, a crucial step in bone regeneration. Consistent with the Masson’s trichrome findings, HE staining (Figure 8B) further corroborates the superior osteogenic response elicited by ZnMF@PEEK. The HE stained images reveal more pronounced cortical bone formation in the vicinity of the osseointegration interface in the ZnMF@PEEK group compared to the PEEK group.

**Figure 8.**
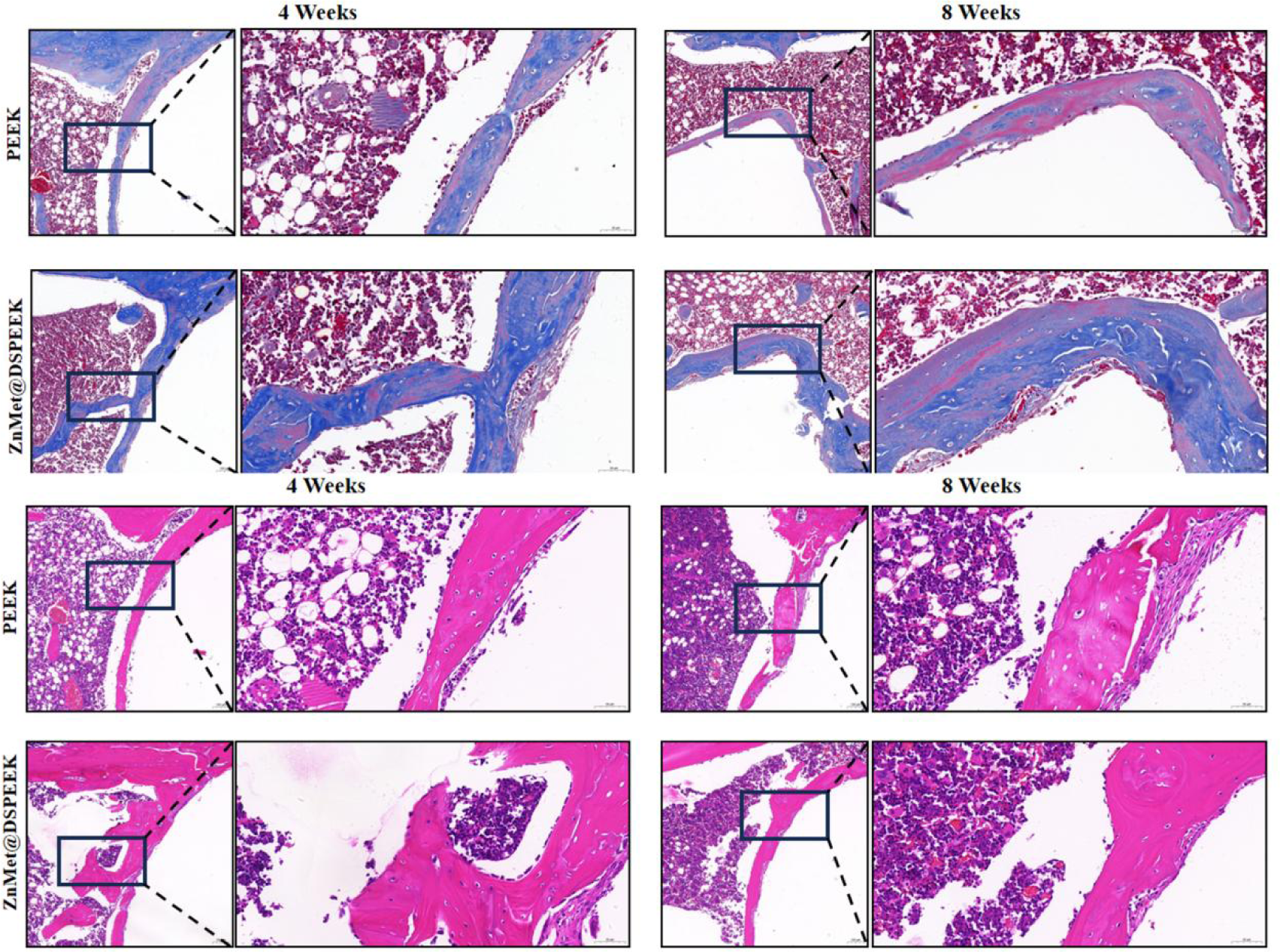
Histological Evaluation of Osteointegration. (A) Masson staining shows higher collagen fiber distribution (blue) and improved organization in ZnMF@PEEK vs. PEEK. (B) HE staining reveals increased viable cells at the osseointegration interface in ZnMF@PEEK compared to PEEK.

To investigate the potential ameliorative effects of ZnMF@PEEK on cellular senescence, a phenomenon that is induced by inflammation and oxidative stress following tissue trauma, which subsequently impacts tissue regeneration efficiency more prominently in elderly individuals due to their inherently reduced regenerative capacity, we conducted immunohistochemical staining for P53, P21, and combined hematoxylin and eosin (HE) with β-gal staining on tissue sections. The results are presented in Figure 9. Figure 9A illustrates the P53 staining outcomes. Notably, there were no significant differences in P53 expression between the PEEK group and the ZnMF@PEEK group at both the 4th and 8th weeks post-operation. This suggests that ZnMF@PEEK does not significantly alter P53 expression levels, a key regulator of cellular senescence and stress response. Figure 9B displays the P21 staining results. Interestingly, a reduced expression of P21 was observed in the ZnMF@PEEK group compared to the PEEK group at both the 4th and 8th weeks. P21, a cyclin-dependent kinase inhibitor, plays a crucial role in cell cycle arrest and senescence. Its decreased expression in the ZnMF@PEEK group indicates a potential attenuation of cellular senescence processes. Lastly, Figure 9C presents the combined HE and β-gal staining results. At the 8th week, a discernible difference emerged, with the ZnMF@PEEK group exhibiting lesser β- gal expression. β-gal, a marker of cellular senescence, was detected in blue-stained cells within normal bone tissue in the PEEK group, suggesting the presence of senescent cells potentially associated with aging. Similarly, blue-stained cells were observed at the interface between the PEEK material and bone tissue, indicating the presence of senescent cells in this region. However, in the ZnMF@PEEK group, while blue-stained cells were still present within the bone tissue, they were noticeably absent at the interface between the material and bone, implying a reduction in senescent cells in this critical area. These findings suggest that ZnMF@PEEK may exert a positive effect on mitigating cellular senescence in the vicinity of implanted materials, particularly at the material-bone interface, which could ultimately enhance tissue regeneration efficiency.

**Figure 9:**
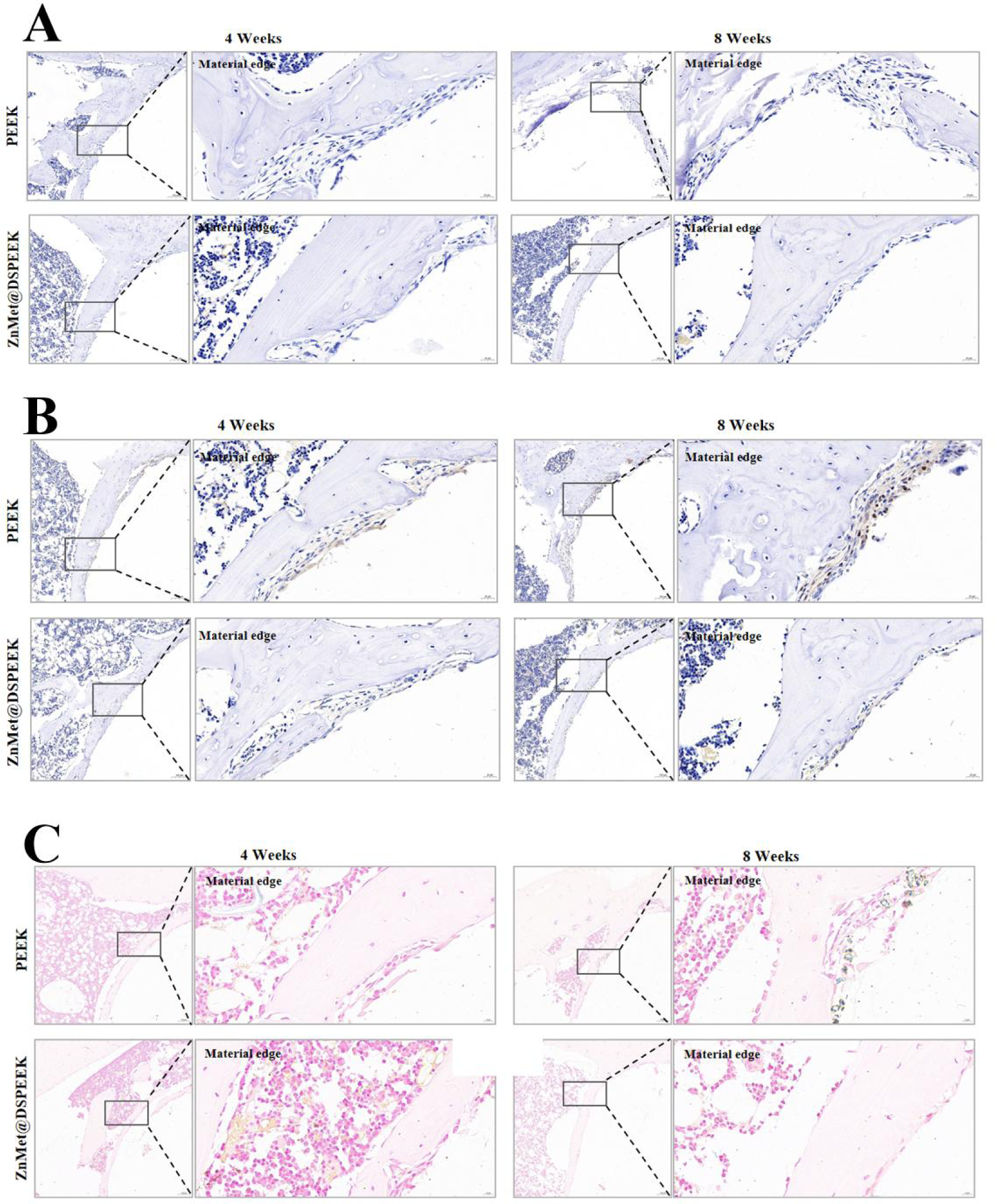
Evaluation of ZnMF@PEEK Effects on Cellular Senescence with Immunohistochemistry of Cell Senescence-Related Proteins, Including (A) P53, (B) P21, and (C) β-gal Staining (Combined with HE)

## 4. Discussions

### 4.1 ZnMF@PEEK was successfully fabricated as promising dental implant

The successful preparation of ZnMF was evident from the SEM (Figure 1A and 1B) and UV-Vis absorption spectra (Figure 2A) analyses. The SEM images revealed a significant transformation from the regular prismatic crystal structure of MF (Figure 1A) to a disordered granular structure in ZnMF (Figure 1B), indicating the formation of a new complex. This structural change was further corroborated by the UV-Vis spectra, which showed an enhancement in the UV absorption properties of ZnMF compared to MF. The more pronounced and higher absorbance intensity peak centered around 255 nm suggested an alteration in the electronic configuration and/or increased conjugation within the ZnMF molecule upon zinc complexation[23, 24].

The successful fabrication of ZnMF@PEEK was confirmed through SEM (Figure 1E and 1F), EDS (Figure 1G-J), and FTIR (Figure 2B) analyses. SEM images demonstrated that ZnMF@PEEK maintained a similar microstructure to DSPEEK but with a slight reduction in pore size and distinctive granular objects evenly dispersed on its surface (Figure 1F). EDS analysis further confirmed the presence of zinc on the surface of ZnMF@PEEK (Figure 1J), indicating the successful incorporation of ZnMF onto the DSPEEK surface. FTIR spectra of ZnMF@PEEK revealed a novel peak at 3364 cm^-1^, attributed to N-H symmetric stretching vibrations, arising from the introduction of ZnMF (Figure 2B)[28].

Interestingly, the wettability characteristics of the samples showed an unusual increase in the contact angle of SPEEK compared to pure PEEK, despite the introduction of hydrophilic sulfonic acid groups upon sulfonation (Figure 2C and 2D)[29]. This anomaly could be attributed to the continued acid etching of the PEEK surface during sulfonation, which might have led to the formation of a porous structure with inherently hydrophobic properties, as reported previously[27]. The further decrease in wettability observed in DSPEEK and ZnMF@PEEK should be due to the hydrophilic nature of dopamine and ZnMF. ZnMF@PEEK, in particular, demonstrated the smallest contact angle among all tested samples, indicating the most enhanced wettability (Figure 2C and 2D). This improved wettability, coupled with the porous structure, could have potential benefits for subsequent osseointegration by facilitating better cell-implant interaction and facilitating mass exchange with the surrounding biological environment[27, 37].

The Zn ion release profile of ZnMF@PEEK over a 24-hour period showed a rapid release within the first 8 hours, followed by a gradual slowing down and reaching a plateau phase after approximately 12 hours (Figure 3). This release pattern can be attributed to our unique sample preparation process, in which ZnMF solution was directly dropped onto DSPEEK followed by freeze-drying. The rapid release phase is likely due to the release of ZnMF that was physically adhered within the porous structure of the sample, relying only on Van der Waals forces rather than strong adhesion to the dopamine. This preparation method can overcome the binding limits of commonly-used dopamine-assisted PEEK surface modified method, as the porous structure on SPEEK was fully used to store more bioactive substances. The rapid release of Zn ions in the initial phase could be beneficial in addressing the acute inflammatory response following the implantation[38–40].

In summary, the existing experimental data fully support the successful synthesis of ZnMF@PEEK. The porous structure, high wettability, and controlled release of bioactive substances exhibited by ZnMF@PEEK suggest that it could offer significant advantages in dental implant applications, and so demonstrate its potential as an advanced dental implant material.

### 4.2 ZnMF@PEEK exhibits superior osteogenic and anti-inflammatory capabilities in *vitro*

The enhanced cell viability and low toxicity observed for ZnMF@PEEK(Figure 4), specifically at the optimal ZnMF concentration of 500 μg/ml, stem from the complementary actions of zinc’s potent anti-inflammatory properties and metformin’s anti-seffects[41–44]. This unique blend fosters a hospitable cellular microenvironment that significantly promotes the growth and proliferation of MC3T3-E1 osteoblasts, pivotal cells for osseointegration.

The remarkable upregulation of RUNX2 and COL1A genes observed in MC3T3-E1 cells cultured on ZnMF@PEEK surfaces (Figure 5A and 5B) underscores the material’s exceptional osteogenic potential. The incorporation of ZnMF on PEEK, likely synergistically enhances the osteogenic signaling pathways. Zn, a known stimulator of osteoblast differentiation and mineralization, should contribute to the upregulation of RUNX2, a master regulator of osteogenesis[45–48]. Meanwhile, MF, with its reported anti-senescence and potentially osteogenic effects, may further potentiate this process[48, 49]. The concurrent increase in COL1A expression, a key structural component of bone matrix, suggests that ZnMF@PEEK not only initiates the osteogenic cascade but also facilitates the actual formation of bone matrix. The significant suppression of inflammatory gene expression, particularly IL-1β (Figure 5C) and IL-6 (Figure 5D), by ZnMF@PEEK highlights its potential to mitigate post-operative inflammation, a common complication in dental implant surgeries. The anti-inflammatory properties of Zn and MF, individually and in combination, likely contribute to this effect. Zn has been shown to reduce pro-inflammatory cytokine production, while MF is known to exhibit pleiotropic effects, including anti-inflammatory actions[50, 51]. By inhibiting inflammation, ZnMF@PEEK may reduce implant rejection rates, accelerate healing, and enhance patient comfort. The combination of osteogenesis and anti-inflammation positions ZnMF@PEEK as a promising material for dental implants.

The downregulation of COL1A in the PEEK group compared to the control, coupled with the upregulation in DSPEEK, suggests that material surface modifications might play a pivotal role in regulating osteogenic gene expression. The sulfonation process used to prepare SPEEK likely introduced hydrophilic sulfonic acid groups, which may have contributed to the anti-inflammatory effects observed[27]. Additionally, the microporous structure created during sulfonation could have facilitated cell adhesion, migration, and differentiation, leading to the upregulation of RUNX2 and COL1A[52].The significantly lower expression of IL-1β in both the PEEK and DSPEEK groups compared to the control suggests that the modifications applied to the PEEK surface have an inherent anti-inflammatory effect. This reduction in pro-inflammatory cytokine expression can be attributed to several factors, including the introduction of sulfonic acid groups during sulfonation and the influence of surface topology on cellular gene expression[13, 53]. The progressive increase in RUNX2 expression from control to PEEK to DSPEEK groups indicates that surface modifications positively regulate osteogenic gene expression and reduce inflammation. The enhanced osteogenic differentiation potential of ZnMF@PEEK is evident from both the cellular and molecular levels. Immunofluorescence staining for alkaline phosphatase (ALP) (Figure 6A and 6B) revealed significantly higher expression levels in the ZnMF@PEEK group compared to the controls and other modified PEEK surfaces. The concerted increase in both ALP activity and osteogenic gene expression underscores the ability of ZnMF@PEEK to stimulate osteoblast differentiation. Moreover, the superior bone mineralization capacity of ZnMF@PEEK, as evidenced by Alizarin Red staining (Figure 6C), highlights its potential to promote bone matrix formation. The superior osteogenic and bone mineralization capabilities of ZnMF@PEEK can be attributed to the synergistic effects of zinc and MF on the PEEK surface.

### 4.3 ZnMF@PEEK exhibit superior osteointegration performance and biocompatibility in *vivo*

The remarkable bone integration capabilities demonstrated by ZnMF@PEEK in the in *vivo* femoral defect model (Figure 7) stem from its multifaceted effects at the molecular and cellular levels. The upregulation of osteogenic genes like RUNX2 and COL1A observed in*vitro*translates to enhanced bone formation in *vivo*, as evidenced by the micro-CT images showing increased new bone formation around ZnMF@PEEK implants compared to PEEK, particularly at 8 weeks post-implantation. The significantly higher BMD and BV/TV values obtained from quantitative CT scan analyses (Figure 7B) further confirm ZnMF@PEEK’s superior osteointegration abilities. Additionally, histological evaluations (Figure 8) reveal a higher distribution of collagen fibers and more pronounced cortical bone formation in ZnMF@PEEK samples, indicative of improved bone matrix organization and osseointegration. Collectively, these findings highlight the synergistic effects of zinc and MF in stimulating osteogenic differentiation, facilitating bone matrix production, and reducing inflammation at the implant site, ultimately leading to the superior bone integration observed with ZnMF@PEEK[41–46].

The potential ameliorative impact of ZnMF@PEEK on cellular senescence, as evidenced by immunohistochemical staining results (Figure 9), underscores its distinctive capacity to augment bone integration in the context of aging populations. Specifically, the decreased expression of senescence markers P21 (Figure 9B) and β-gal (Figure 9C) at the implant-bone interface in ZnMF@PEEK-treated specimens indicates a potential mitigation of cellular senescence. This effect can be attributed to the combined anti- senescence mechanisms inherent in ZnMF@PEEK. Zinc’s potent anti-inflammatory properties indirectly contribute to combating cellular senescence by suppressing the inflammatory microenvironment that exacerbates senescent phenotypes[54]. Concurrently, metformin exerts direct anti-senescence effects by reducing P21 expression, thereby potentiating the overall anti-senescence capability of ZnMF@PEEK. The synergistic interplay of these mechanisms enables ZnMF@PEEK to effectively counteract cellular senescence[55], fostering more efficient bone healing and integration in elderly patients, with implications for improved outcomes in dental implant applications targeting the geriatric demographic.

The lack of discernible tissue damage in critical organs, as evidenced by the H&E staining outcomes (Figure 10), underscores the clinical safety of ZnMF@PEEK. Elderly patients often face challenges related to reduced osteogenic potential, persistent inflammation, and increased cellular senescence, all of which can compromise implant success[56]. ZnMF@PEEK addresses these challenges by stimulating bone formation, mitigating inflammation, and combating cellular senescence. The favorable biocompatibility of ZnMF@PEEK, coupled with its superior bone integration capabilities and anti-senescence effects, positions ZnMF@PEEK as a promising material for dental implant surgery in the elderly population.

**Figure 10.**
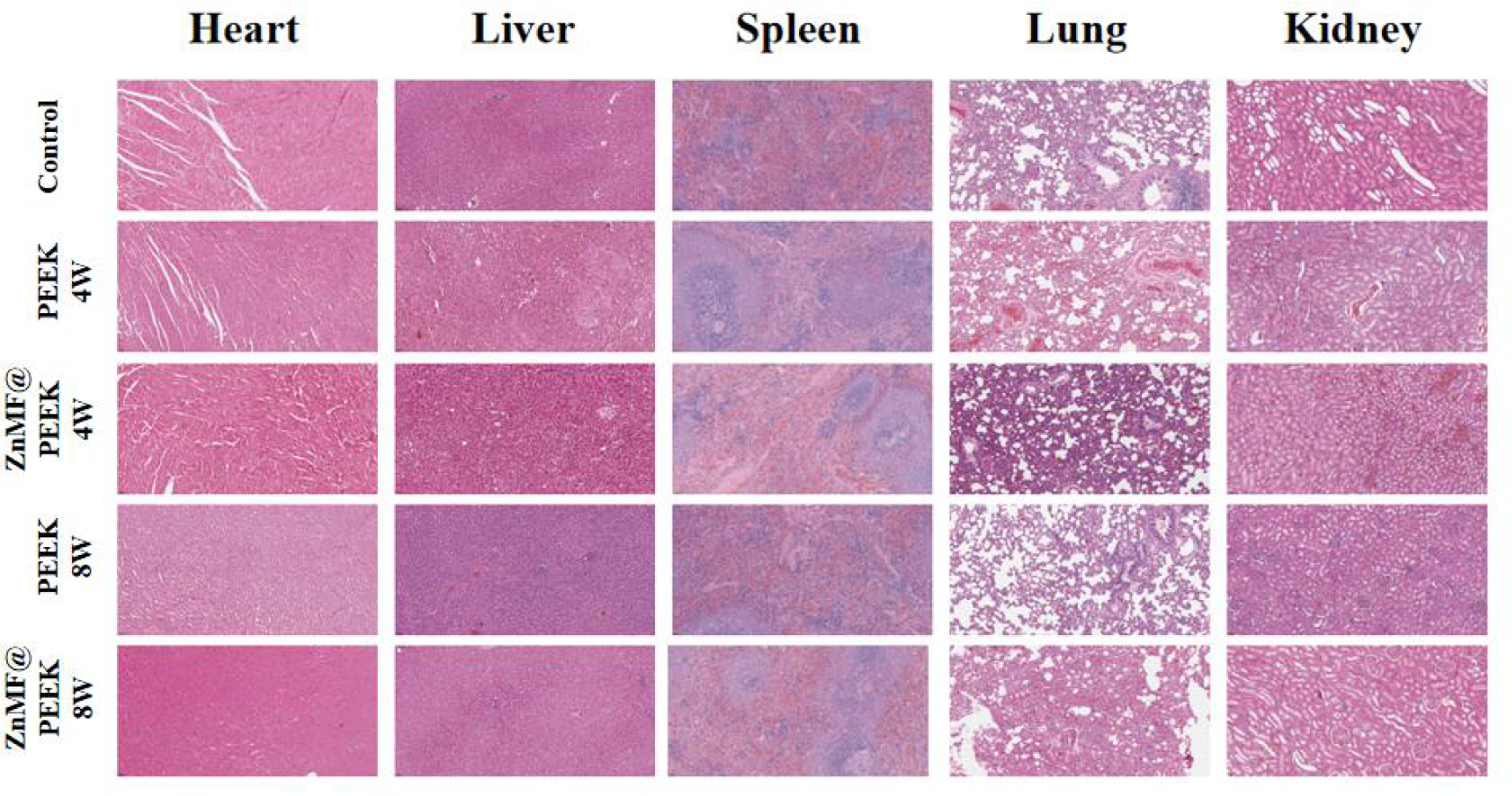
presents the H&E staining outcomes of vital organs following 4 and 8weeks after surgery, with implantation of PEEK and ZnMF@PEEK. Notably, no discernible tissue damage was observed in the critical organs across all groups, encompassing the heart, liver, spleen, lung, and kidney. This observation suggests that the ZnMF@PEEK sample we developed should be clinical safety in *vivo*.

## Conclusions

In this study, we successfully developed a new functionalized PEEK derivative, ZnMF@PEEK, with tailored bioactivities including anti-inflammatory, osteogenic, and anti-senescence properties, as a dental implant for the elderly population. Physicochemical characterization confirmed the successful synthesis of ZnMF@PEEK, and in *vitro* experiments further demonstrated its remarkable multifunctional bioactivity, particularly its osteogenic and anti-inflammatory properties. Specifically, ZnMF@PEEK promotes osteoblast differentiation and bone mineralization while effectively attenuating inflammatory responses and mitigating cellular senescence. These findings were further validated through therapeutic evaluation using a rat femoral defect model, which exhibited enhanced bone formation at the implant-bone interface, increased bone mineral density, and reduced cellular senescence. Importantly, ZnMF@PEEK demonstrated excellent biocompatibility, with no significant tissue damage to critical organs, thereby ensuring its clinical safety. Therefore, we believe that ZnMF@PEEK should be a promising dental implant material for the elderly, offering a novel solution to the multifaceted challenges of osseointegration.

